# PENGUIN: A rapid and efficient image preprocessing tool for multiplexed spatial proteomics

**DOI:** 10.1101/2024.07.01.601513

**Authors:** A. M. Sequeira, M. E. Ijsselsteijn, M. Rocha, Noel F.C.C. de Miranda

## Abstract

Multiplex spatial proteomic methodologies can provide a unique perspective on the molecular and cellular composition of complex biological systems. Several challenges are associated to the analysis of imaging data, in particular regarding the normalization of signal-to-noise ratios across images and background noise subtraction. However, straightforward and user-friendly solutions for denoising multiplex imaging data that are applicable to large datasets are still lacking. We have developed PENGUIN –Percentile Normalization GUI Image deNoising: a rapid and efficient image preprocessing tool for multiplexed spatial proteomics. In comparison to existing approaches, PENGUIN stands out by eliminating the need for manual annotation or machine learning model training. It effectively preserves signal intensity differences and reduces noise, thereby enhancing downstream tasks like cell segmentation and phenotyping. PENGUIN’s simplicity, speed, and user-friendly interface, deployed both as script and as a Jupyter notebook, facilitate parameter testing and image processing. We illustrate the effectiveness of PENGUIN by comparing it with conventional image processing techniques and solutions tailored for multiplex imaging data. This comparison underscores PENGUIN’s capability to produce high-quality imaging data efficiently and consistently.

## Introduction

In recent years, the field of multiplexed imaging technologies has grown tremendously, allowing for spatially resolved proteomic, transcriptomic, or metabolic profiling of biological systems [1, 2]. Imaging Mass Cytometry (IMC) [3] and Multiplex Ion Beam Imaging (MIBI) [4] make use of metal-conjugated antibodies for the detection of proteins by means of mass spectrometry. This process involves quantifying isotopic reporter masses released from the tissue following ablation of small tissue areas with a laser beam or ion beams. Standard immunofluorescence (IF) [5] and high dimensional fluorescence methods, like Co-Detection by IndEXing (CODEX) [6], make use of fluorescently labeled antibodies.

Despite the increasing application of multidimensional proteomics spatial technologies, their resulting data poses challenges that often cannot be overcome with conventional image analysis approaches. A common issue is the presence of noise in data, which must be removed prior to analysis [7]. Noise sources can vary depending on which antibodies, detection channels and tissue types are used and can be manifested in artifacts such as hot pixels or background noise [7-9]. Furthermore, some protein markers may exhibit weak signal and low signal-to-noise ratios [10]. Together, these noise sources deteriorate the image quality and complicate downstream analyses of multiplex imaging data. Therefore, robust and stable methods for denoising have become increasingly important with the application of these methods [7].

In this work, we propose an improved pipeline for denoising multiplexed proteomics - PENGUIN - Percentile Normalization GUI Image deNoising. It uses scaling, thresholding, and percentile-based filters to tackle different sources of noise in multiplex images. We demonstrate its applicability by using an existing IMC dataset and benchmark it against existing methods for image preprocessing, as well as methods designed specifically for analysis of multiplex imaging data. Furthermore, by also applying PENGUIN to multiplex IF images, we demonstrate that this method can be widely applied to different types of imaging data. The tool and the benchmarking data are available on https://github.com/deMirandaLab/PENGUIN.

### Current approaches for multiplex imaging normalization

#### Noise and denoising strategies

Image denoising plays a crucial role in computer vision, yet it remains a challenging task since it is essential to reduce noise without losing relevant signals [11]. Noise can follow various probabilistic distributions, with Gaussian noise - where noise pixel values follow a Gaussian distribution - and impulse noise, such as salt-and-pepper noise, being the most common. In impulse noise, the noise is random and unrelated to the image pixels. Typically, in a clean image, neighboring pixels are correlated, as objects or surfaces often span multiple adjacent pixels, whereas noise in neighboring pixels is usually uncorrelated. Thus, a basic and effective denoising strategy applies a predefined operation within a local neighborhood – like a 3×3 pixel window – to produce an output value for the center pixel, creating a processed image as the filter is applied across each pixel in the input image. Filters, which are quick and straightforward to use, can significantly enhance image preprocessing and noise removal. They can be linear, such as Gaussian and mean filters, or non-linear, like the median filter [11, 12]. Linear filtering normalizes pixels by replacing them with the mean value of their neighborhood or, in the case of Gaussian filters, a weighted mean based on the proximity of neighboring pixels. While this approach reduces outlier pixels, it can also blur edges and significantly diminish high-frequency image details. Moreover, with this approach, impulse noise is averaged out but is not eliminated entirely [12, 13].

On the other end, non-linear filters operate without linear operations. The most prevalent among these are order-statistic filters, which include median filters. In this approach, the center pixel is substituted with the median of the ranked values from its surrounding pixels. Median filters excel in dealing with impulse noise, as such noise usually ranks at the extreme ends of the brightness scale. Additionally, median filtering not only significantly diminishes the spikes caused by impulse noise, but also maintains the brightness differences across signal transitions. This leads to minimal blurring of regional edges, thus preserving the integrity of boundary positions within the image [12]. However, median filters are primarily effective against impulse noise, requiring that the noisy pixels constitute less than half of the neighborhood area, and their performance declines when noise is present in more than 20% of an image’s pixels. They are less successful in removing Gaussian or random-intensity noise, as the brightness of noisy pixels does not significantly differ from that of their neighboring pixels [12]. Percentile filters, akin to median filters, adjust pixel values based on a range of percentiles rather than solely the median (50^th^ percentile). Offering similar advantages to median filters, percentile filters have the same benefits as the median filters and offer flexibility as different markers may benefit from different values of noise reduction, as they may display more or less shot noise [12].

Other non-linear filters include Bilateral, Non-Local Means, Adaptive Filtering, Total Variation Denoising, Anisotropic Diffusion, and Wavelet denoising filters.

Recently, deep learning (DL) has become a powerful tool in image analysis, introducing innovative strategies for denoising. These strategies can be categorized in two types: those that are trained using ground truth data, and those that do not require it. In many application areas, such as spatial omics, obtaining clean ground truth images can be challenging. Thus, denoising methods that operate independently of a ground truth dataset are particularly valuable. Presently, Noise2Void stands out as a widely utilized algorithm in this category [14]. It trains a neural network to eliminate noise by leveraging information from the surrounding pixels within the same image, obviating the need for a separate, clean reference image. Another method involves denoising autoencoders. An autoencoder is a neural network employed in unsupervised learning, aimed at learning efficient data representations by compressing the data into a compact internal form, and then reconstructing the original data from this representation. When used for denoising, the autoencoder is trained to process noisy input data and output clean data, thereby learning to ignore the noise during the reconstruction of the original data. This training process enhances the autoencoder’s ability to identify relevant features in the presence of noise, making it a valuable tool for various image restoration and denoising tasks [15].

#### Denoising imaging mass cytometry (IMC) data

With the advent of multiplex spatial phenotyping technologies, specialized approaches to address image noise, like crosstalk and hot pixels, have been developed. Crosstalk, a phenomenon where signals from one channel interfere with adjacent channels, can lead to the misidentification of signals, especially for co-expressed markers [7, 10]. To mitigate this, methods like CATALYST [9] employ pre-acquisition signal compensation matrices, while post-acquisition solutions are also available [8, 16]. However, carefully designed antibody panels can often minimize the need for correction by reducing low-intensity channel overlap [10, 17].

Hot pixels, another common artifact in IMC images, are identified as individual pixels exhibiting significantly higher signal intensities compared to their surroundings and are typically modeled as a Poisson process. This randomness, characterized by the independent occurrence of points in a mathematical space, can be attributed to detector abnormalities and can be influenced by various factors, including differences in metal isotope conjugation, antibody concentration, and spatial arrangement [10, 17]. Additionally, small clusters of consecutive hot pixels, not representative of biological structures may form due to nonspecific antibody binding, antibody aggregates, or contamination by dust particles [8, 10].

Hot pixels are often dealt with by applying traditional image filtering methods and thresholds. Researchers have devised numerous “homebrew” computational strategies, tailored specifically to individual projects [16, 18, 19]. A notable method, featured in Steinbock, a Python toolkit for processing multiplexed tissue images, addresses hot pixels by comparing each pixel’s value to those of its eight surrounding neighbors. Should the difference between the pixel and any of its neighbors surpass a predetermined threshold, the pixel’s value is reduced to match that of the highest neighboring pixel value [20].

In addition to conventional denoising techniques, tools like Ilastik offer pixel classification in a supervised manner, enabling the distinction between background noise and true signal pixels for each marker [20-22]. Essentially, an experienced user trains the Ilastik Random Forest pixel classifier by manually identifying pixels as either representing signal or background. This trained model is then applied across all the images, resulting in a binary expression map, where non-noise pixels are assigned a value of 1.This method has proven effective in not only eliminating background noise, but also in standardizing and normalizing signals across different samples, thereby mitigating batch effects [22]. However, the requirement for manual annotations across a series of images within a cohort can be labor-intensive. This manual curation process is particularly challenging when dealing with large datasets, making it a less desirable approach in such contexts.

Recently, Lu et al. [17] introduced IMC-Denoise, a two-step pipeline combining a Differential Intensity Map-based Restoration (DIMR) algorithm for removing hot pixels and a self-supervised DL algorithm for shot noise image filtering, inspired by Noise2Void, named DeepSNiF. Initially, hot pixels are identified using Anscombe transformation differential maps. Subsequently, DeepSNiF processes these images by randomly masking pixels through a stratified sampling approach. DL approaches, however, tend to be time consuming, resource heavy and require rigorous tuning of parameters.

Despite the availability of various IMC denoising approaches, there is still no consensus on the optimal approach, and the absence of a standardized strategy hinders dataset comparisons. Current techniques either necessitate manual marking of noisy pixels across all markers – a process that is both time-consuming and prone to inconsistency – or fail to adequately address all noise types and nonspecific antibody signals.

In this study, we propose that the primary IMC noise sources, such as hot pixels and background noise, can be effectively managed using straightforward classical algorithms. PENGUIN is quick, scalable, reproducible, and does not rely on manual pixel annotation nor requires extensive hardware resources.

## Results

To mitigate noise in IMC images, the authors identified several key considerations:

1. Shot noise, characterized by random pixels exhibiting significantly higher signal intensities compared to their surroundings (also known as salt and pepper noise), can be efficiently addressed by filtering out very sparse signals, namely pixels that lack neighboring signals.
2. Non-specific antibody binding, when it occurs, results in signal areas that are less intense than those from specific binding.
3. Noise tends to occur independently across different channels within the same images, meaning it can affect one channel without necessarily impacting others. Nonetheless, within a specific cohort, noise generally displays consistent patterns across all images from the same channel.

To address noise in IMC images, the authors put forward a method that integrates scaling, thresholding, and percentile-based filters. First, images undergo saturation at the 99th percentile to remove the impact of extremely bright pixels. Following this, signal intensities across each channel and image are normalized to a scale from 0 to 1. A thresholding technique is then applied to discard background signals, essentially removing signals of low intensity. Subsequently, hot pixels are pinpointed and eliminated using percentile filters. A crucial element of this strategy is the use of the percentile filter exclusively for identifying and removing pixels without altering the image’s clarity or edges. So, after post-processing, the image retains its sharpness and edge integrity. Furthermore, the normalization of values from 0 to 1 facilitates their direct application in subsequent analyses, like cell segmentation. The foundational principle behind PENGUIN, the percentile-based noise detection, is illustrated in Figure 1.

**Figure 1.**
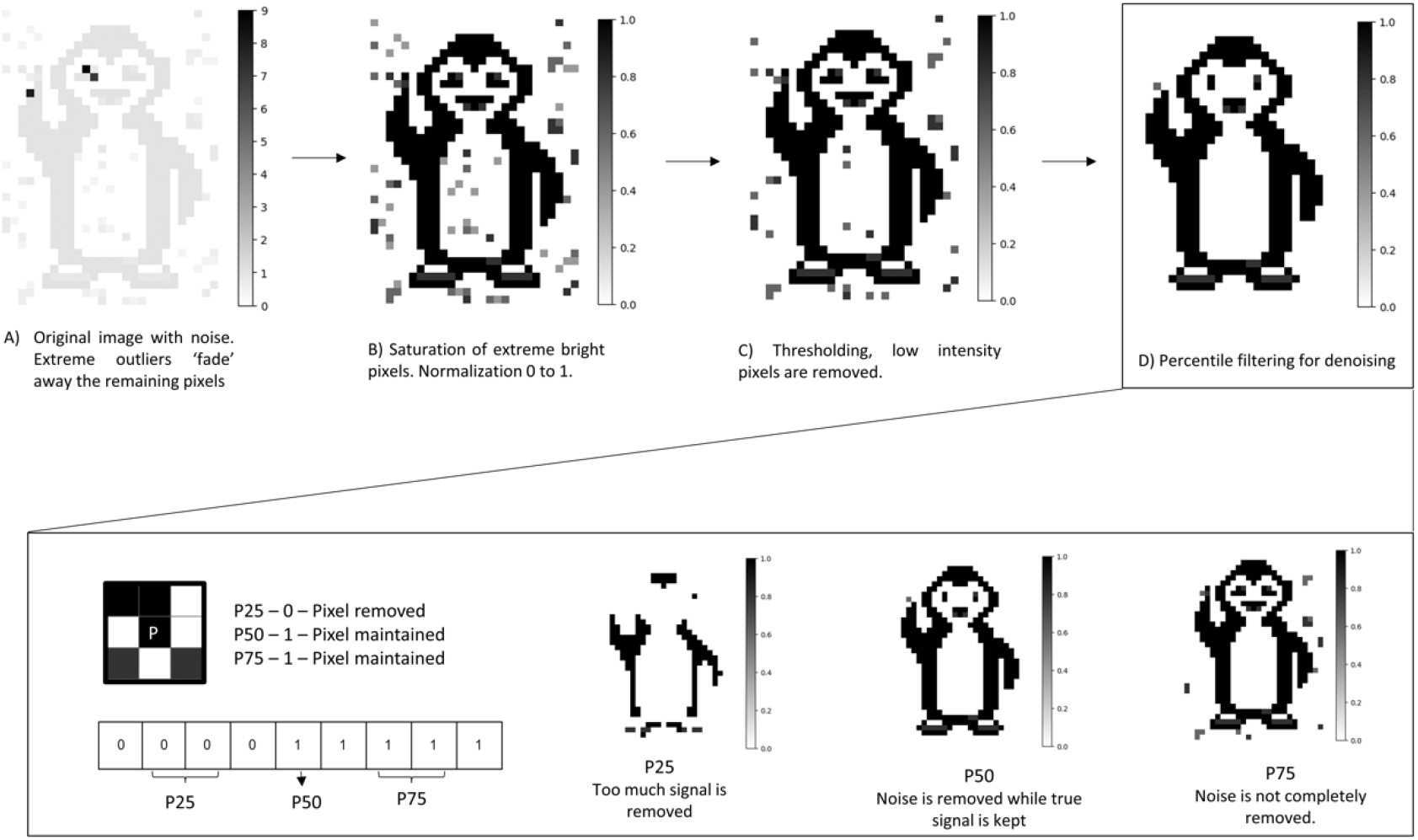
Overview of the PENGUIN pipeline. A) The original image contains noise and extreme outliers, which obscure the rest of the signal. B) Outliers are saturated by setting values above the 99^th^ percentile to this value, enhancing visualization. Each channel and image are then normalized between 0 and 1. C) Pixels with values below a chosen Threshold are removed. D) Percentile filtering is applied to detect and remove noise pixels. For each central pixel, a 3×3 window is considered. Pixels are transformed to Boolean (0/1) and ordered ascendingly. If the percentile value is 0, the pixel is considered noise and set to 0; if it is above 0, it is considered a real signal. For example, a 50^th^ percentile (P50) requires that 5 out of 9 pixels to be positive, the 75^th^ percentile requires 2 out of 9 pixels to be positive, and the 25^th^ percentile (P25) requires 7 out of 9 pixels to be positive. Depending on the channel characteristics, low percentile values may lead to the removal of true signals, while high percentile values may retain noise signals.

Key adjustable parameters in this pipeline include the Threshold (T) value and the Percentile (P) value, which are both critical to its performance. A higher T will result in the removal of more signal in the initial phase of the pipeline. If the output of a channel is predominantly background signal with only high intensity values representing true signals, the T should be set high. In contrast, for channels with minimal background noise, the T can be lower or even omitted. The P parameter plays a crucial role in the pipeline, with lower values like 25 being stricter as they result in the removal of nearly all sparse signals, while higher values such as 75 are more lenient, allowing for more sparse signals. Tailoring these parameters for each channel is essential due to significant differences in the behavior of markers.

Figure 2 illustrates the effects of varying T and P parameters on the IMC imaging data of β-catenin, CD20, Vimentin, CD45, and FOXP3. For example, β-catenin showed optimal noise reduction at a setting of T0.1 and P50, whereas CD20 required a higher threshold of T0.3 to effectively eliminate noise. On the other hand, for Vimentin and CD45, clear images were achieved at P25, while a T setting was not required.To simplify the visualization and deployment of the preprocessing pipeline, we created a user-friendly Jupyter notebook. This tool enables direct observation of how thresholds and percentiles affect different markers to help decide which thresholds are more applicable to each marker and image. A sneak peek of the notebook can be seen in Figure 3.

**Figure 2.**
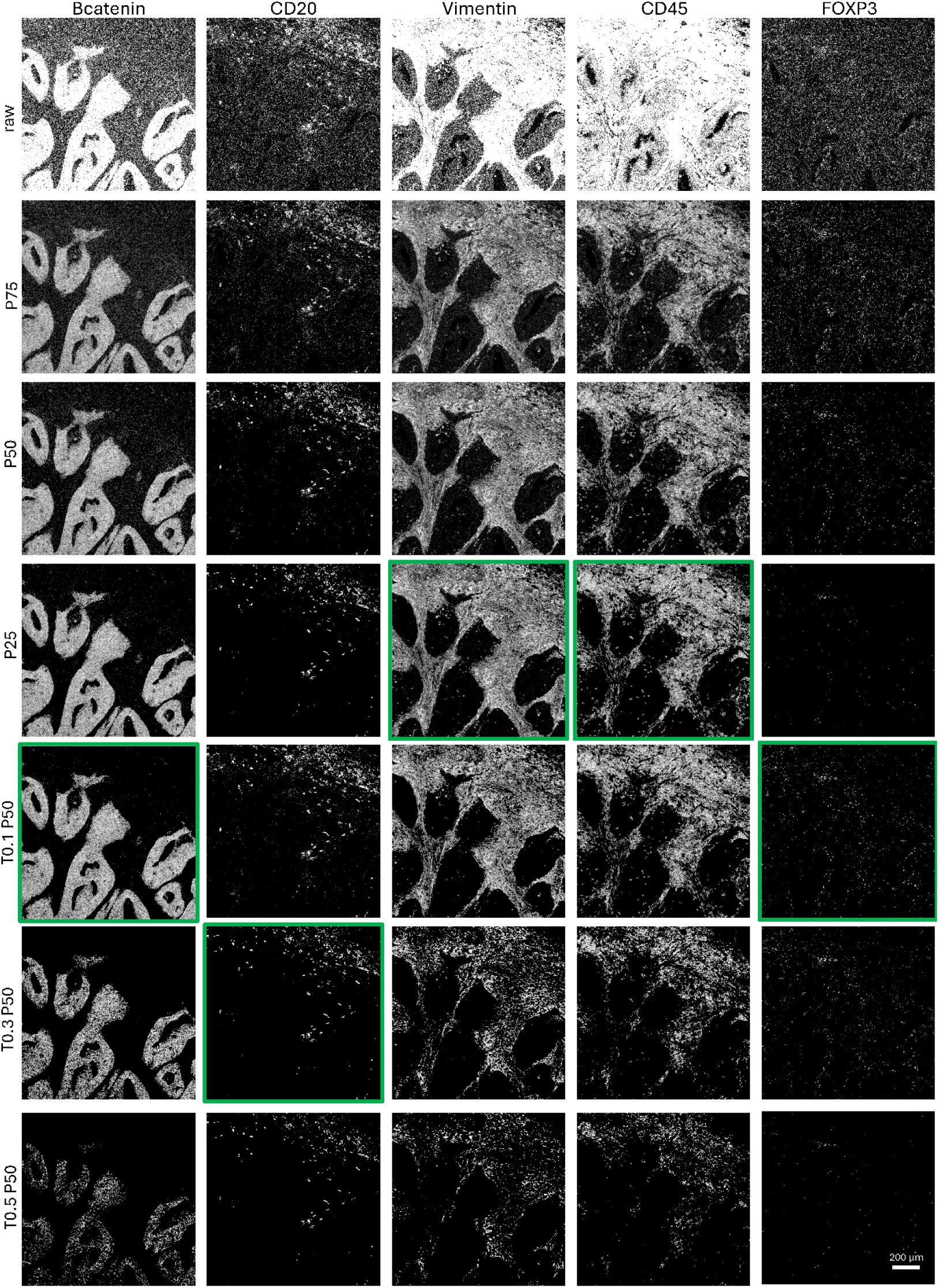
Application of different percentiles (P90,P75,P50,P25,P10) and thresholds (T0.1,T0.3,T0.5, with P50) for β-catenin, CD20, Vimentin, CD45 and FOXP3. The optimal parameters for each channel are highlighted in green.

**Figure 3.**
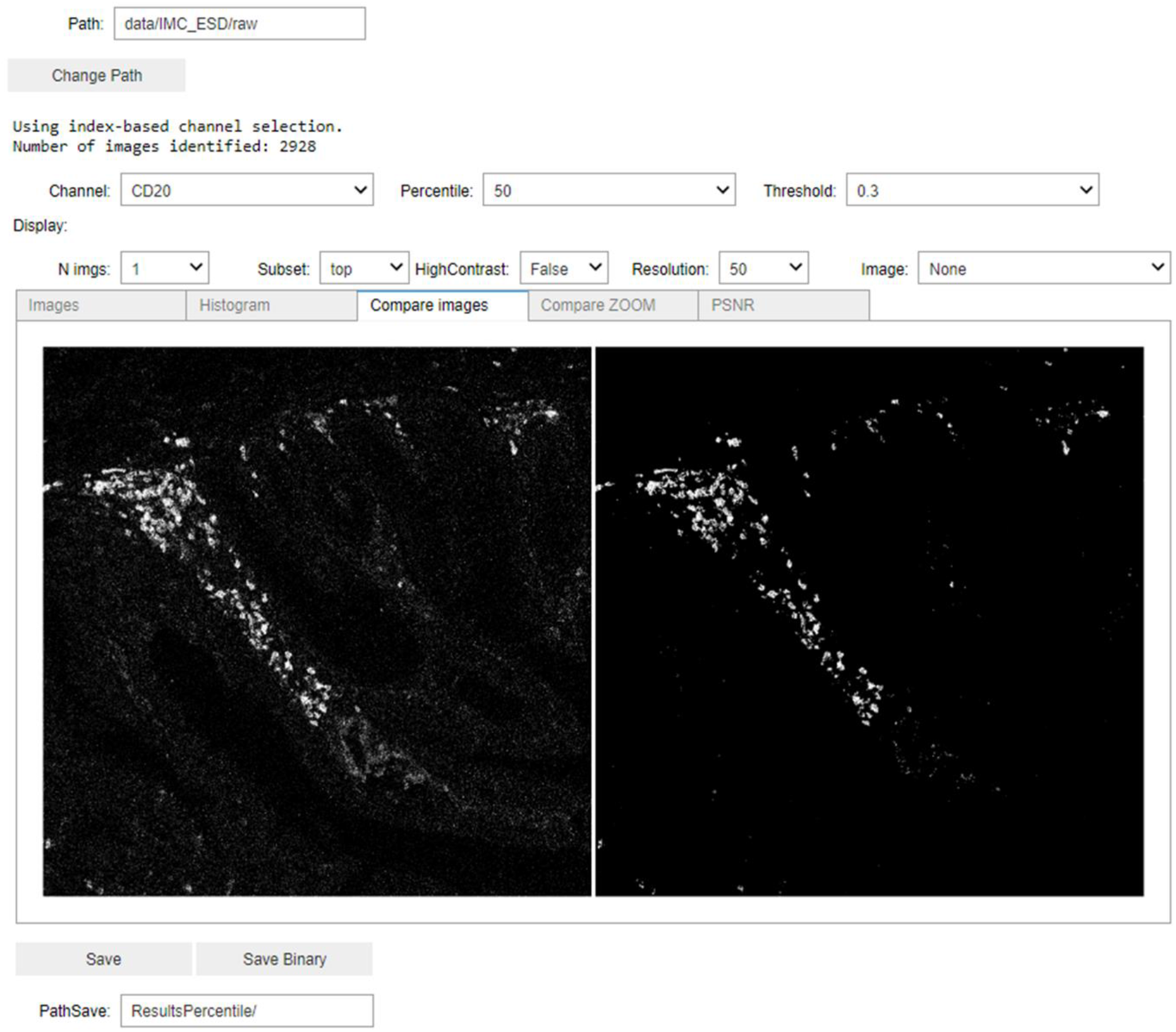
PENGUIN Jupyter Notebook sneak peek

The code for the preprocessing pipeline is freely available. The users can apply the functions for direct scripting or use the notebook provided for a more user-friendly experience.

### Comparison of several methods for denoising IMC

PENGUIN was evaluated against both traditional image denoising techniques and IMC-specific approaches using a public dataset comprising 39 cellular markers evaluated by IMC on 61 samples [24].

To improve the performance of standard image denoising methods, we applied saturation to the 99^th^ percentile to remove bright outliers and normalized each channel’s signal intensity between 0 and 1 [22]. We initially explored various classical filtering methods, including Gaussian, mean, percentile, non-local means, bilateral, total variation, wavelet, anisotropic diffusion, and BM3D filters (see supplementary figures 1 & 2). As anticipated, linear filters such as Gaussian and mean filters tended to blur images, leading to a loss of detail and definition in tissue boundaries (supplementary figures 1 & 2, linear filters). Although these filters did not directly eliminate hot pixels, the pixels became less conspicuous as they were blurred. The remaining classical image filters either failed to adequately reduce noise (for example anisotropic and bilateral filters), or distorted the images, like the non-local mean filter, rendering them unsuitable for this type of data.

We also evaluated DL methods designed for denoising images without the need for paired ground truth data, specifically Noise2Void [14] and denoising autoencoders [23], which were trained for 20 or 50 epochs. These approaches did not effectively denoise the images and led to image distortion, suggesting they may not be applicable for IMC images. However, it is important to note that DL models perform better with larger datasets. As the availability of IMC image datasets increases, the performance of these models may improve. It is also worth mentioning that our experiments only involved a basic convolutional denoising autoencoder configuration, and the potential for optimization through different combinations of layers, epochs, and parameters is vast. Yet, DL methods require significant time for development, tuning, and training, and often necessitate specialized hardware that may not be widely accessible.

Next, we applied two IMC-specific methodologies. Firstly, we explored a hot pixel filter [20] that uses a 3×3 kernel to assess if the difference between a pixel and its maximum neighboring value exceeds a predefined threshold; if so, the pixel is classified as noise and is attributed the maximum neighbor’s value. This method was applied to both normalized, using thresholds ranging from 0.05 to 0.2, and to raw images with a default threshold of 50. Although this approach succeeded in removing some hot pixels, the processed images still contained noise, and occasionally, real signals were lost (supplementary figures 1 & 2, IMC specific strategies). Secondly, with IMC Denoise [17] we applied both the DIMR algorithm alone and in combination with the DeepSNIF DL algorithm, adhering to the authors’ guidelines and default settings. The DIMR algorithm alone proved inadequate for removing hot pixels. When supplemented with DeepSNIF, similar to the issues observed with Noise2Void, some image distortions occurred. As with other DL methods, the potential configurations for IMC Denoise are virtually limitless; however, fine-tuning the correct parameters is a labor-intensive process. Additionally, distinct models must be trained for each channel, further complicating the application of this approach.

Finally, we applied the methodology developed at our lab [22], where we developed a semi-automated background removal (SABR) pipeline based on Ilastik (Figure 4). Compared to the raw data, SABR effectively removes most noise, producing clear binary images. However, this process also results in some loss of specific signals, noticeable for instance in the case of β-catenin immunodetection (Figures 4 and 5).

**Figure 4.**
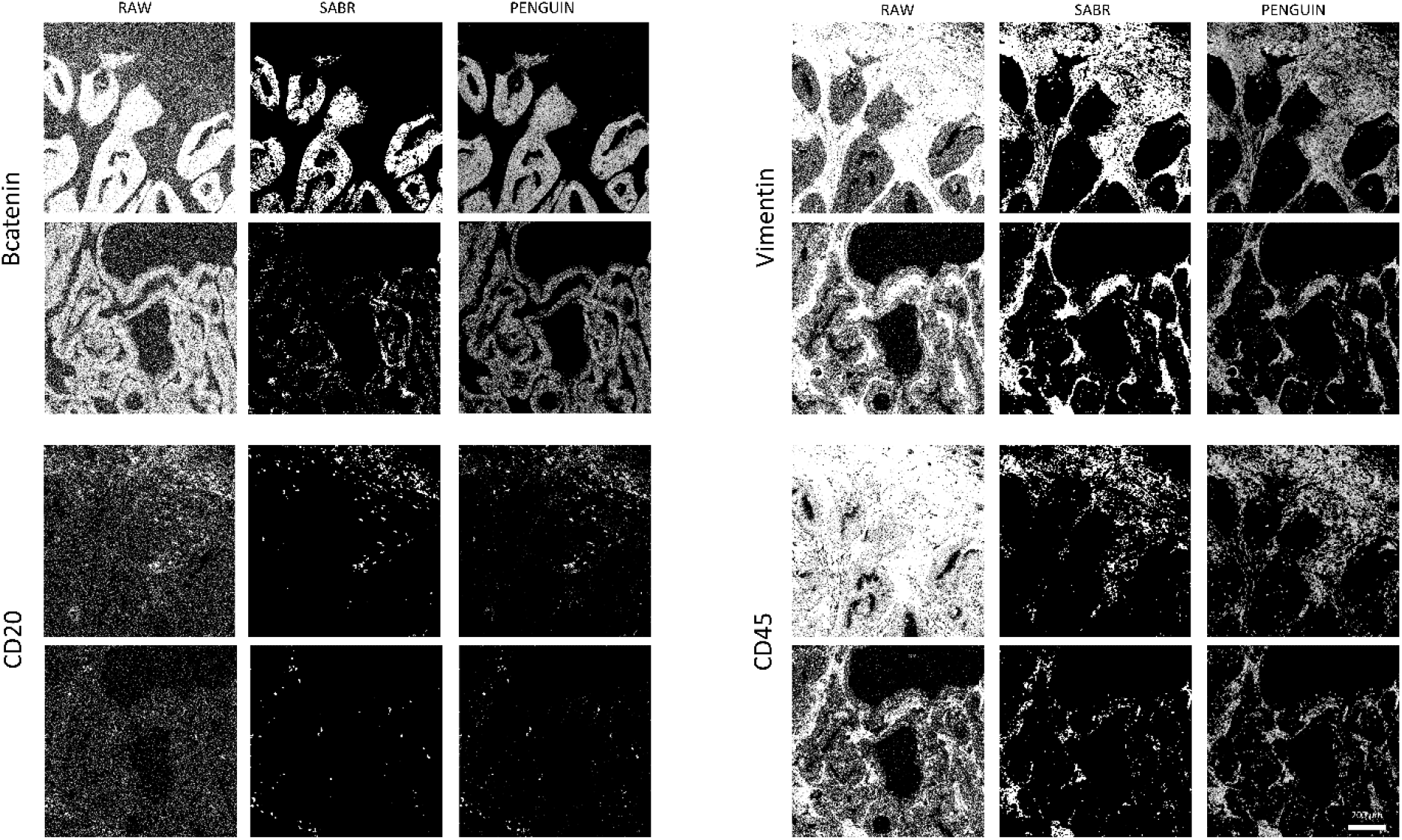
Bcatenin, CD20, Vimentin and CD45 markers for two regions of interest with raw images, PENGUIN processed and Ilastik-based SABR approach.

**Figure 5.**
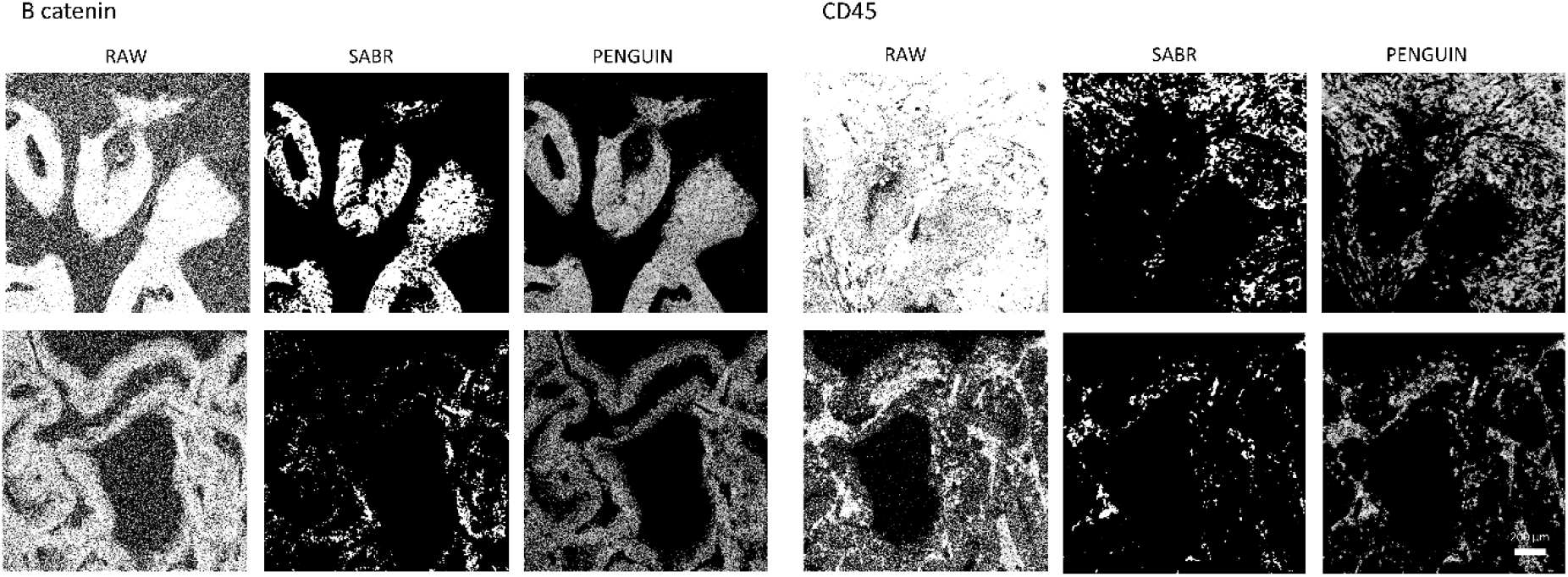
Zoom in from figure 4 for B catenin and CD45 markers. When compared to the SABR approach [22]], PENGUIN retains more real signal and the key aspects of cell structure.

When applying PENGUIN to the same set of images, we found that it effectively removes noise from raw images, matching the quality achieved by the Ilastik-based SABR method. Additionally, PENGUIN often retains more signal than SABR, as highlighted in Figure 5.

This underscores the effectiveness of PENGUIN in eliminating noise from IMC images. There are also substantial differences between PENGUIN and the SABR pipeline. Firstly, PENGUIN necessitates manual adjustment of thresholds and percentiles for each channel but does not require the development of specific classifiers to distinguish between signal and noise for each marker. Secondly, unlike the Ilastik-based method, which generates binary images indicating merely the presence or absence of signals, PENGUIN produces images with normalized values ranging from 0 to 1, representing various signal intensities. This allows for a more nuanced analysis, as PENGUIN not only retains more signal, but also preserves critical details of cellular structures.

Setting the thresholds and percentiles is an intuitive process, made even simpler and more efficient with the use of the provided notebook. Processing all 61 images for each channel took less than 4 seconds on an Intel(R) Core(TM) i7-9700 CPU, which represents a significant time saving compared to training and deploying Ilastik models for each marker. Additionally, the results are both fully reproducible and easily achievable, marking a significant improvement over existing methods.

### PENGUIN enhances downstream analysis

After confirming the effectiveness of PENGUIN for image processing, we evaluated its impact on downstream analysis. This involved comparing raw, unprocessed images with those corrected using the Ilastik-based SABR method and the PENGUIN pipeline in terms of cell segmentation and cell phenotyping/clustering.

A significant advantage of PENGUIN over the previous approach is that its output can be directly utilized in cell segmentation pipelines, eliminating the need for training an additional Ilastik model to differentiate between membrane, nucleus, and background markers, thereby saving considerable time. Moreover, as highlighted earlier and depicted in Figure 5, PENGUIN’s normalization process retains more signal and crucial cell structural features.

To assess the efficacy of these methods, we combined the normalized images with their corresponding cell segmentation masks to calculate relative marker expression per cell. Cells were then clustered using t-SNE to identify cell subsets, as shown in Figure 6. Consistent with findings by Ijsselsteijn et al. [22], normalization and background removal are essential for accurately identifying signals and cells, as well as normalizing IMC inter-sample variation for automated downstream analysis. Similar to the current state-of-the-art techniques, PENGUIN adjusts for interpatient variability, achieving well-defined cells and clusters. This represents an improvement over existing methods because it is faster, reproducible, and allows for the preprocessed images from PENGUIN to be directly used in cell clustering and phenotyping, bypassing the necessity for two rounds of training Ilastik models.

**Figure 6.**
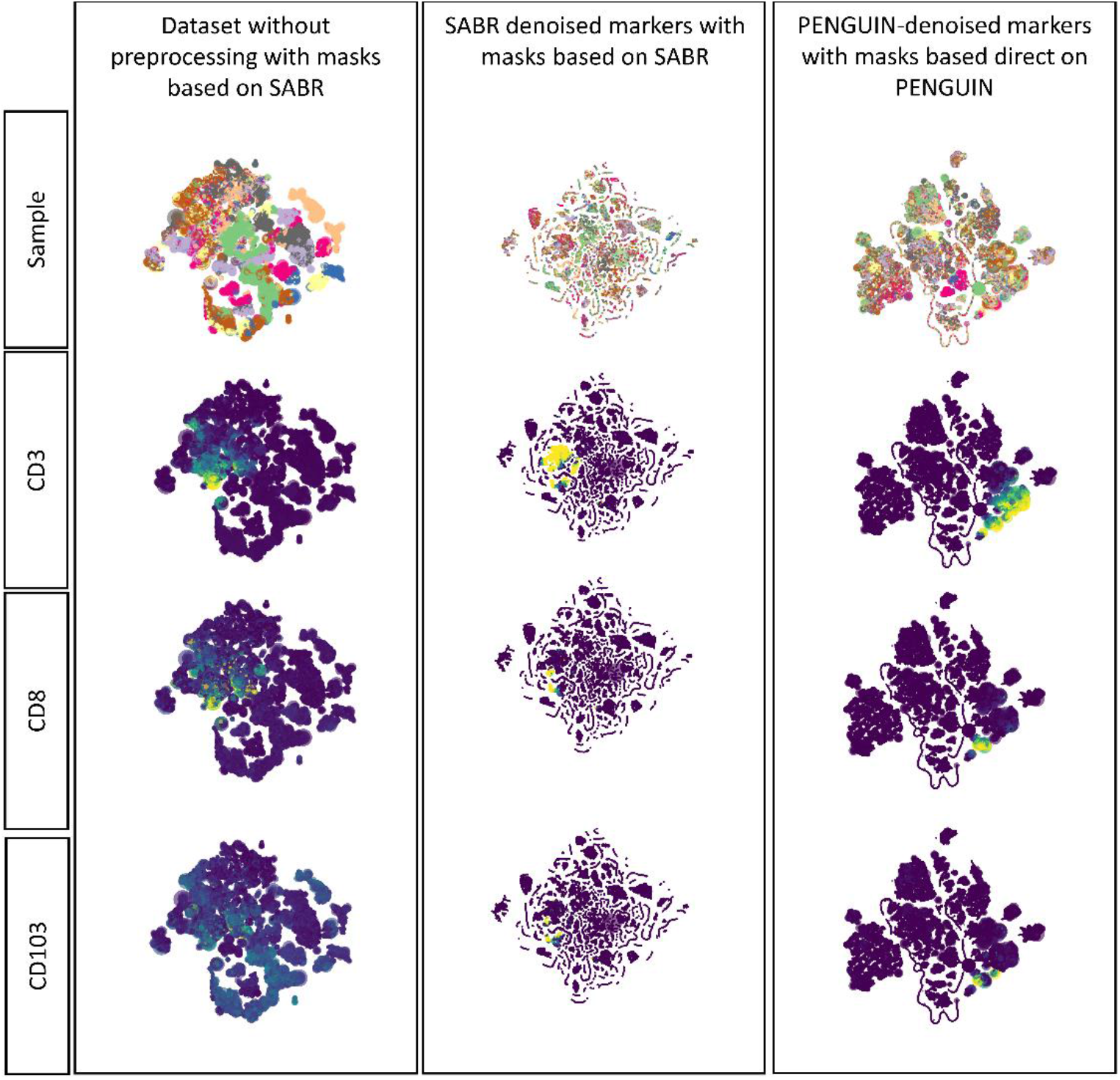
t-SNE obtained by A) raw marker expression and SABR Ilastik-based cell masks, B) SABR Ilastik-denoise marker expression and SABR Ilastik-based cell masks, C) PENGUIN-denoise marker expression and PENGUIN-based cell masks. The t-SNE are colored by sample, CD3, CD8 and CD103 markers.

### PENGUIN can be applied to other image modalities

Various imaging modalities in multiplex spatial proteomics, such as MIBI, CODEX, and multiplex IF, produce different types of images. Encouraged by the promising outcomes achieved with the PENGUIN pipeline, and given that the signal characteristics from these modalities share similar limitations, the authors extended the application of their developed pipeline to a set of in-house generated IF images. The results, showcased in Figure 7, reveal high-quality denoised images, achieved with significantly reduced time, resources, and operator-induced variations. Consequently, PENGUIN facilitates the normalization of both IMC and IF images for downstream analysis without compromising the specificity of the antibody signal. Additionally, its speed and scalability render it highly effective for processing large and expanding imaging datasets.

**Figure 7.**
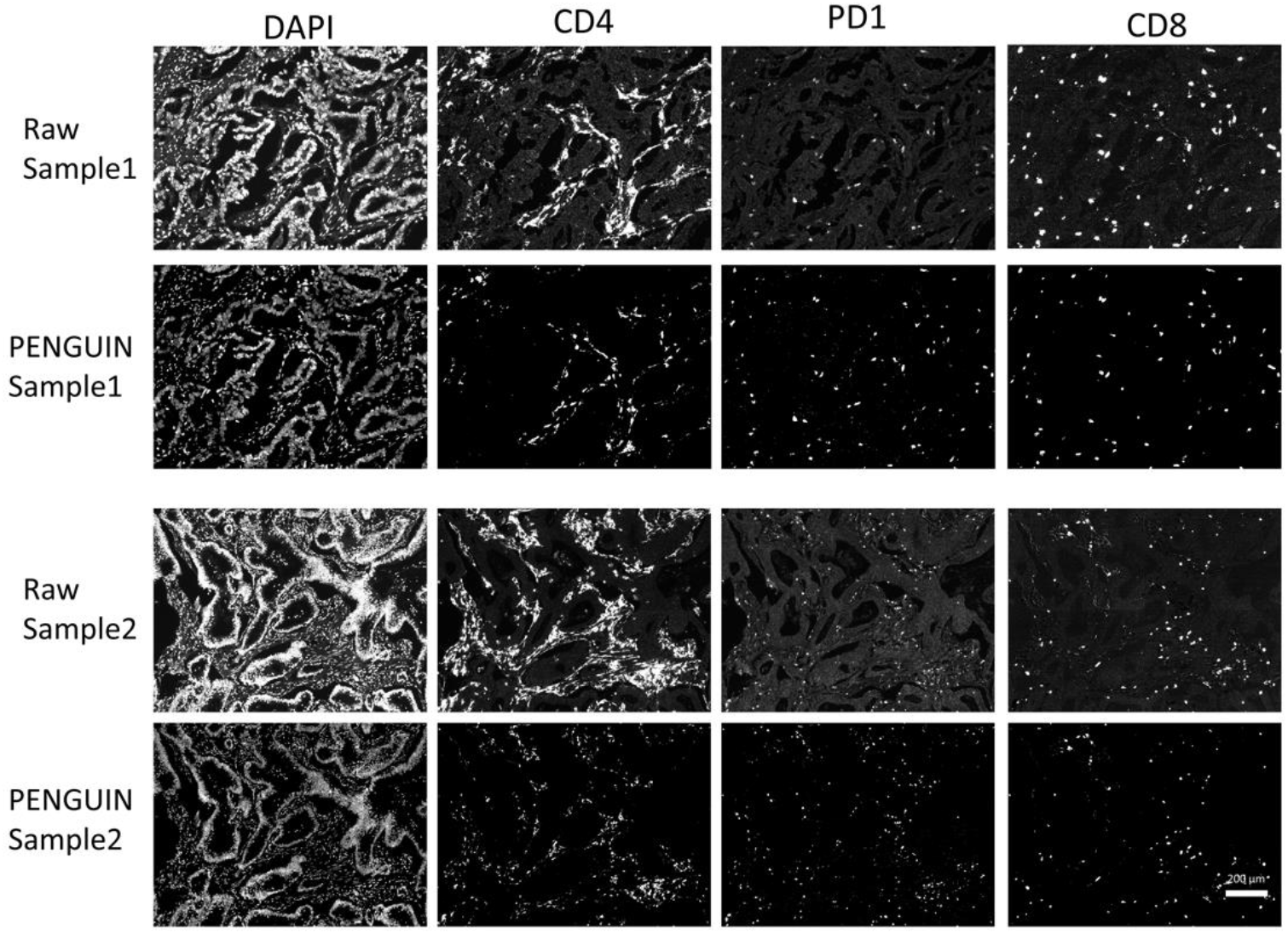
PENGUIN applied to IF images, namely, DAPI, CD4, PD1 and CD8 signal.

## Discussion

Spatial proteomics data, like that from IMC, provides a distinctive insight into the spatial organization of tissues, offering broad applications. However, handling signal-to-noise ratios, particularly with extensive antibody panels, poses significant challenges. Computational preprocessing methods are essential for ensuring the reliability of subsequent analyses [7]. Although various custom algorithms have previously achieved success in denoising IMC data, there is a growing need for a standardized, efficient preprocessing pipeline to manage the increasing volume of data. Given that pixel intensity values in spatial proteomics vary across targets, and signals for low-abundance antigens are often sparse, it is logical that the density of positive pixels provides more information about the presence of a signal than the intensity of individual pixels.

In this context, we introduce PENGUIN, which effectively denoises IMC images through scaling, thresholding, and percentile filters, yielding excellent results. We evaluated this pipeline against traditional filtering methods, DL-based approaches, and IMC-specific solutions, demonstrating that PENGUIN delivers clear images quickly, easily, and reproducibly. A Jupyter notebook with a user-friendly interface is provided, facilitating the testing of various thresholds and percentiles per marker and simplifying the process of adjusting parameters, transforming, and saving images. Additionally, the core code is available for scripting.

Compared to the recently developed Ilastik-based SABR method, PENGUIN eliminates the need for manual annotation or training of machine learning models, preserves signal intensity differences across tissues, reduces noise, and retains more structural details. Moreover, PENGUIN enhances essential downstream tasks such as cell segmentation and phenotyping. Due to its simplicity, speed, intuitiveness, high-quality results, reproducibility, and ease of deployment, we believe PENGUIN is a valuable tool to aid in the analysis of IMC data and, eventually, of other multiplex imaging platforms.

## Methods

All the tests were run in a Intel(R) Core(TM) i7-9700 CPU. The DL models were run using a GPU Nvidia TU102 GeForce RTX 2080 Ti.

### Dataset

All the results for IMC were obtained using the publicly available cohort of 61 IMC raw images with 39 markers of colorectal cancer [24].

### Denoising Methods comparison

Before application of the denoising filters, images were saturated to the level of the 99th percentile and normalized between 0 and 1. Using raw images was also tested, however, normalizing leading to significant improvements in image quality.

The following filters were applied to each channel: Gaussian filter using sigma values of 0.7 and 1; mean filter; percentile filter (with values of 25,50 and 75); non-local means (patch size 5, patch distance 11 and sigma 0.2); bilateral filter; total variation (defined as proposed by Chambolle, weight 0.3); Wavelet; anisotropic diffusion and BM3D filtering(sigma 0.1, 0.2 and estimated sigma that retrieve values between these two values were tested). Parameters not specified were set as default. Gaussian, mean and percentile filters were based on SciPy implementation [25]. Non-local means, bilateral filter, total variation and wavelet filtering were based on scikit-image package [26]. Bm3D was retrieved from source package [27].

The filter to remove hot pixels in IMC images was implemented as described by Windager et al. [20] which applies a Scipy maximum filter with size 3*3. If the difference between the center pixel and its maximum neighbor is above a defined threshold is set to the value of the maximum neighbor. Thresholds ranging from 0.05 to 0.2 were applied with normalized images, default threshold, 50, was used with raw images.

Noise2Void was trained as defined by Krull et al. [14] per channel with all images of the cohort with 0.2 of data used for validation. Loss was defined as MSE neighborhood radius 5 and UNET kernel size 3 for a patch size of 64. Models were trained for 20, 50, and 100 epochs.

Convolutional autoencoder for image denoising were built as defined in [23] with the encoder being defined as 2 consecutive 2D convolution layers (32 filters kernel size (3,3), activation relu) and max pooling (pool size (2,2) and the decoder as 2 Transpose convolutional layers (32 filters kernel size (3,3), activation relu) and a last Convolutional operation with 1 filter, and sigmoid activation. Adam optimizer and binary cross entropy were used. The training process was done for 20 or 50 epochs per channel. The DL implementation was using Tensorflow [28]. All the parameters not specified were set as default.

IMC Denoise [17] was implemented following the instructions from the tutorial notebook and default parameters were defined. IMC Denoise has two different steps, the DIMR algorithm and the DeepSNIF. We tested only applying the DIMR algorithm per channel (4 neighbors, window size3 and 3 iterations) and DIMR algorithm followed by the DeepSNIF trained for 200 epochs.

### PENGUIN implementation

PENGUIN is implemented in Python 3.10 and deployed as a code package and Jupyter Notebook. For each image and each channel, pixels above the 99th percentile are set to that value and are normalized from 0 to 1. After this, pixels with values below a defined threshold are removed and a percentile filter is used to detect which pixels are considered noise or signal (window 3*3). The same thresholds and percentiles per channel are applied to all the images in the cohort. To demonstrate the concept we applied the percentiles of 90, 75, 50, 25, and 10 and percentile 50 after thresholding pixels below 0.1, 0.3, and 0.5. This comparison was done for Bcatenin, CD20, Vimentin, CD45, and FOXP3 for their importance as markers and different signal distributions. There are two Jupyter Notebooks, one for the case where each region of interest has one TIFF file per channel, and the other where there is a TIFF stack containing all channels.

### Comparison of methods for downstream analysis

#### PENGUIN parameters definition

The raw images were analyzed by an IMC expert, using PENGUIN notebook for better visualization. Parameters, namely thresholds and percentiles, were set for each channel using the Notebook tool (**Erro! A origem da referência não foi encontrada**.) and applied to all images in the cohort.

#### Ilastik-based background removal

For each marker, a random forest classifier was trained in Ilastik [21] to recognize background pixels and signal pixels, as described by Ijsselsteijn et al. [22]. After rigorous training on approximately 10% of the dataset, the random forest classifier was applied to all images of a given marker and data was exported as simple segmentation masks where all background pixels were set to zero and signal pixels to one.

#### Cell segmentation

Cell masks were created using CellProfiler V3.0 [29]. For comparison purposes, masks were created with raw data, as well as with PENGUIN normalized images. Using raw data, the traditional Ilastik/CellProfiler combination was used, where first probability masks for nucleus, membrane and background were generated in Ilastik using the DNA, Vimentin and Keratin images. Next, these were loaded into a CellProfiler pipeline to identify primary objects (nuclei) after which membranes were appended to those using the identify secondary objects module. Visual inspection was done to compare the masks with the original IMC Images. After PENGUIN normalization, the normalized images could directly be used for the creation of cell segmentation masks in Ilastik using the same approach.

#### Cell type phenotyping/cluster

The 2 different masks from cell segmentation (Ilastik-based and PENGUIN) were loaded into ImaCytE

[30] combined with the raw, Ilastik-based denoised, and PENGUIN-based denoised marker profiles to define relative marker expression per cell. FCS files were produced, and clustering of cells was performed by t-SNE to identify cell subsets, using Cytosplore [31].

#### Immunofluorescence labelling and imaging

Immunofluorescence labelling was performed using the OPAL methodology. In short, deparaffinisation, rehydration, endogenous peroxidase blocking and antigen retrieval with 10 mM citrate buffer (pH 6.0) were performed on four µm thick FFPE tissue sections. Next, sections were blocked using superblock solution (Thermo Fisher Scientific) and incubated for 1 hour with anti-CD4. Opal amplification was performed by consecutive incubations with a polymeric HRP-linker antibody conjugate (Immunologic, The Netherlands) for 10 minutes and a 10 minute incubation with OPAL650 reagent (Akoya Biosciences). The sections were boiled for 15 minutes in 10 mM citrate buffer (pH 6.0). Blocking, primary antibody incubation, followed by HRP and OPAL were then repeated for all antibodies (CD103 – OPAL620, PD1 – OPAL570, GZMB – OPAL520, CD8 – OPAL690). Between each step the sections were washed with PBS-tween. Finally, the sections were incubated with one µM DAPI, washed with PBS (without tween) and mounted using Prolong^®^ Gold Antifade Reagent (Cell Signaling Technologies). The slides were imaged and spectrally unmixed using the Vectra 3 imaging system and the InForm software (Akoya Biosciences) after which the component images were normalised with PENGUIN.

Parameters, thresholds and percentiles, were set for each channel (**Erro! A origem da referência não foi encontrada**.) and applied to all images in the cohort.

## Data and code availability

The IMC data is publicly available from Roelands et al. [24] and is accessible on BioImage Archive (Accession number S-BIAD587, https://www.ebi.ac.uk/biostudies/bioimages/studies/S-BIAD587).

The PENGUIN pipeline is completely available and ready to use at https://github.com/deMirandaLab/PENGUIN. The repository also contains all the code to replicate the methods comparison, along with explainable Jupyter notebooks.

## Acknowledgments

This study was supported by the Portuguese Foundation for Science and Technology-FCT and FSE through PhD fellowship: [2020.0786BD].

