## Supplementary Materials for "PENGUIN: A rapid and efficient image preprocessing tool for multiplexed spatial proteomics"

### Supplementary images

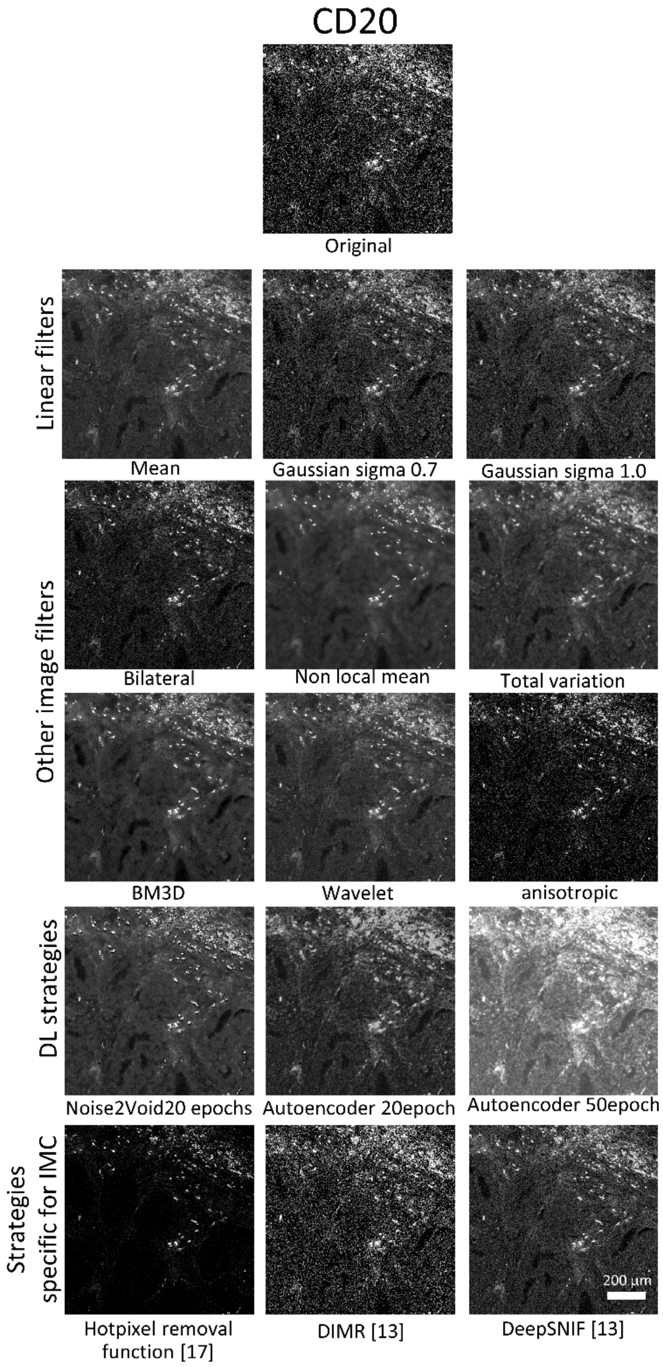


*Supplementary Figure 1 - CD20 channel of one sample with linear filters (mean, gaussian with sigma 0.7 and 1); other traditional image filters ( bilateral, non local mean, total variation, BM3D, wavelet and anisotropic); DL based strategies for denoising, namely Noise2Void and a denoise autoencoder (trained for 20 and 50 epochs) and; Strategies specific designed for IMC, namely the hot pixel filter defined by [20] and DIMR and DIMR with DeepSNIF from [17].*

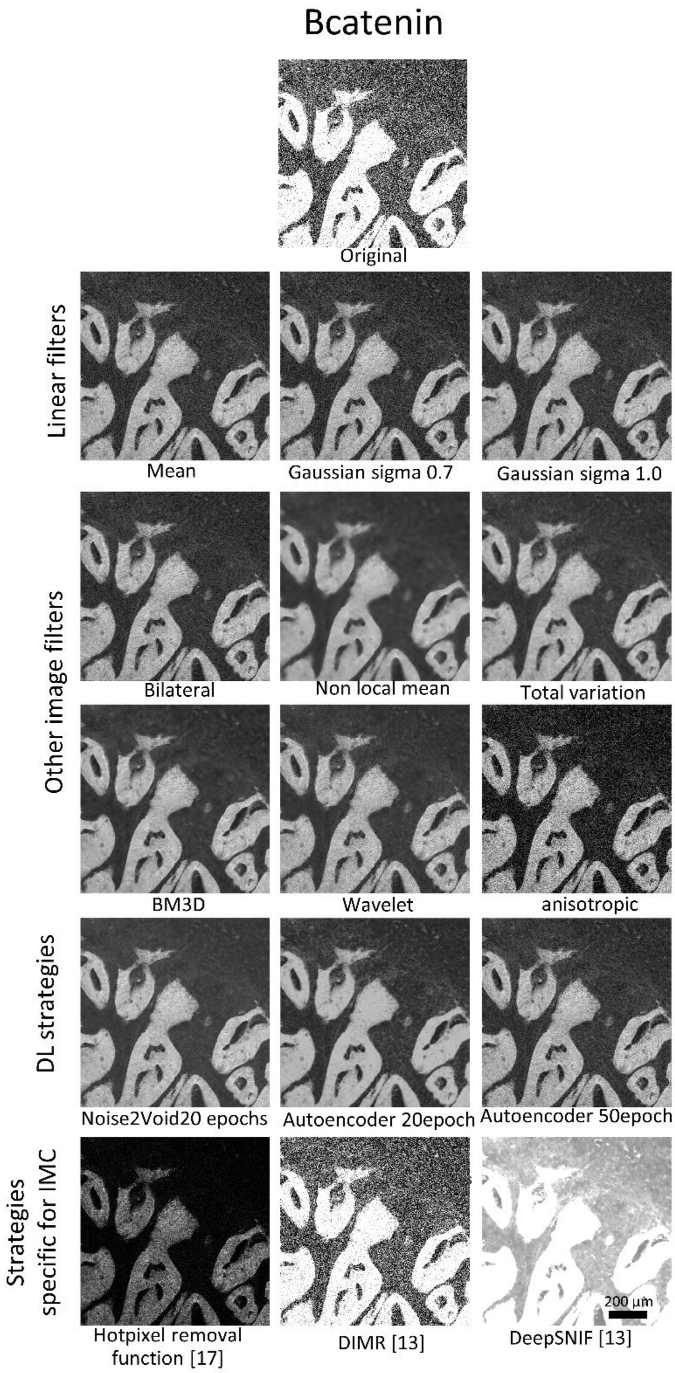


*Supplementary Figure 2 - Bcatenin channel of one sample with linear filters (mean, gaussian with sigma 0.7 and 1); other traditional image filters ( bilateral, non local mean, total variation, BM3D, wavelet and anisotropic); DL based strategies for denoising, namely Noise2Void and a denoise autoencoder (trained for 20 and 50 epochs) and; Strategies specific designed for IMC, namely the hot pixel filter defined by [20] and DIMR and DIMR with DeepSNIF from [17].*

*Supplementary Table 1 - Thresholds and Percentiles values for the 39 IMC markers*

| Marker | Percentile | Threshold | Marker | Percentile | Threshold |
|---|---|---|---|---|---|
| B catenin | 60 | 0.2 | CD8a | 60 | 0.25 |
| CD103 | 30 | 0.1 | cleavedcaspase | 40 | 0.3 |
| CD11b | 50 | 0.25 | D2-40 | 50 | 0.25 |
| CD11c | 40 | 0 | FOXP3 | 50 | 0.1 |
| CD14 | 40 | 0.1 | Granzyme B | 40 | 0.2 |
| CD15 | 40 | 0.25 | HLA-DR | 50 | 0.1 |
| CD163 | 40 | 0.3 | ICOS | 50 | 0.1 |
| CD20 | 50 | 0.1 | IDO | 30 | 0.3 |
| CD204 | 50 | 0.1 | Keratin | 50 | 0.5 |
| CD3 | 50 | 0.1 | Ki-67 | 50 | 0.2 |
| CD31 | 50 | 0,2 | P16ink4a | 50 | 0.2 |
| CD38 | 40 | 0.2 | PD-1 | 40 | 0.2 |
| CD39 | 40 | 0.2 | PD-L1 | 50 | 0.3 |
| CD4 | 50 | 0.2 | Tbet | 60 | 0.1 |
| CD45 | 50 | 0.2 | TCRgd | 40 | 0.1 |
| CD45RO | 50 | 0.2 | TGFbeta | 40 | 0.2 |
| CD56 | 50 | 0.2 | Vimentin | 40 | 0.1 |
| CD57 | 60 | 0.2 | VISTA | 50 | 0.2 |
| CD68 | 40 | 0.2 | CD7 | 50 | 0.2 |

*Supplementary Table 2 - Thresholds and Percentiles values for the IF markers*

| Marker | Channel | Percentile | Threshold |
|---|---|---|---|
| DAPI | DAPI | 60 | 0.2 |
| CD4 | Opal650 | 50 | 0.5 |
| PD1 | Opal570 | 40 | 0.7 |
| CD8 | Opal690 | 40 | 0.7 |
